# Infrared Laser Sampling of Low Volumes Reveals Marker Lipids in Palatine Tonsil Carcinoma via Shotgun Lipidomics

**DOI:** 10.1101/2025.06.05.658016

**Authors:** Leonard Kerkhoff, Manuela Moritz, Dennis Eggert, Anna Worthmann, Joerg Heeren, Henrike Zech, Till S. Clauditz, Waldemar Wilczak, Hartmut Schlüter, Christian S. Betz, Arne Böttcher, Jan Hahn

**Affiliations:** Department of Otorhinolaryngology, University Medical Center Hamburg-Eppendorf, Martinistr. 52, 20246 Hamburg, Germany; Section Mass Spectrometry and Proteomics, Center for Diagnostics, University Medical Center Hamburg-Eppendorf, Martinistr. 52, 20246 Hamburg, Germany; Department of Biochemistry and Molecular Cell Biology, Center for Experimental Medicine, University Medical Center Hamburg-Eppendorf, Martinistr. 52, 20246 Hamburg, Germany; Department of Pathology, Diagnostic Center, University Medical Center Hamburg-Eppendorf, Martinistr. 52, 20246 Hamburg, Germany; Mildred Scheel Cancer Career Center HaTriCS4, University Medical Center Hamburg-Eppendorf, Martinistr. 52, 20246 Hamburg, Germany

**Author notes:** These authors contributed equally and share senior authorship (Arne Böttcher, Jan Hahn). **Corresponding author:** Jan Hahn, Dr., Section Mass Spectrometry and Proteomics, Center for Diagnostics, University Medical Center Hamburg-Eppendorf, Martinistr 52, 20246 Hamburg, Germany.

**Keywords:** nanosecond infrared laser, laser ablation, shotgun lipidomics, lipid marker, OPSCC

## Abstract

Complete surgical resection is essential for oropharyngeal squamous cell carcinoma (OPSCC) therapy, underscoring the need for improved intraoperative margin assessment. To advance *in-vivo* diagnostics of OPSCC, Nanosecond infrared laser (NIRL) tissue sampling combined with shotgun lipidomic analysis reveals lipidome differences between OPSCC tissue and adjacent healthy tissue. Ablations were performed on tonsil squamous cell carcinoma on n=28 samples from 11 patients with an established chamber setup and a subset of n=6 samples from three patients with a custom-made handheld applicator. Welch’s t-test results (p=0.05, two-fold change) revealed a similar OPSCC lipid profile in 7 of 11 patients. Potential tumor lipid markers were identified as consistently significantly increased, despite biological heterogeneity of the samples, underscoring their potential diagnostic value. Tissue ablation with a custom-made handheld applicator coupled to a laser fiber was successful and the lipidomic analysis was consistent to the chamber setup. Although our setup is currently limited by an ablation time exceeding one minute, this study demonstrates the potential of the handheld applicator for future applications such as endoscopy or intraoperative diagnostics.

## Introduction

Alterations in cancer cell lipid metabolism, such as an increase in *de novo* lipogenesis, fatty acid uptake and fatty acid oxidation are characteristic properties of almost every human tumor (Broadfield *et al*, 2021). Lipids that are *de novo* synthesized in tumor cells differ from circulating lipids in the body, resulting in altered lipid composition within these cells. These changes contribute, among other factors, to carcinogenesis, immune escape, proliferation and metastasis (Broadfield *et al*, 2021; Martin-Perez *et al*, 2022; Santos & Schulze, 2012; Snaebjornsson *et al*, 2020; Beloribi-Djefaflia *et al*, 2016; Butler *et al*, 2020). To comprehensively analyze the lipidome, mass spectrometry (MS)-based techniques such as shotgun lipidomics, are widely used, providing detailed insights into lipid metabolism (Han & Gross, 2005).

However, lipids and other small molecules are particularly well suited for online ambient mass spectrometry, which allows the direct measurement of many biomolecules with minimal sample preparation. This approach comes with several downsides, including suppression effects, no quantification and limited identification of only small molecules, but opens new possibilities in numerous fields, particularly in diagnostics and cancer surgery (Cooks *et al*, 2006; Takats *et al*, 2017). The differentiation between healthy and tumor tissue for diagnostic use or intraoperative margin assessment is feasible (Ogrinc *et al*, 2022).

Various approaches for ambient mass spectrometry have been developed each with its advantages and limitations. Desorption electrospray ionization (DESI) utilizes a charged solvent stream to extract and ionize the components of the sample and makes them immediately available for MS (Takáts *et al*, 2004; Ifa & Eberlin, 2016; Woolman *et al*, 2017b). It has proven to be applicable in confined spaces, like in a probe (Chen *et al*, 2013). A further non-destructive method for online ambient mass spectrometry is a handheld “MassSpec Pen”, which extracts tissue surface molecules by a water flow over the sampling probe, reflecting only the lipids on the surface (Zhang et al, 2017). Another approach is called “iKnife” and combines electrosurgical cutting via diathermy with collection of the arising aerosol for subsequent MS, termed rapid evaporative ionization MS (REIMS). Good sensitivity and specificity have been demonstrated in the diagnosis of gynecologic tumors (Schäfer *et al*, 2009; Balog *et al*, 2013; Marcus *et al*, 2022). Integration with a harmonic ultrasound scalpel is also possible (Manoli *et al*, 2021).

Pulsed infrared lasers (IRL) utilizing the concept of desorption by impulsive excitation (DIVE) (Franjic & Miller, 2010) represent another promising approach for tissue sampling. With a wavelength of approximately 2940 nm, they are used to specifically target O-H molecular bonds in the molecules. In the tissue, the pulse energy of the IRL is almost entirely absorbed by these bonds, generating a rapidly emerging tissue aerosol while minimizing heat transfer and preventing damage to the surrounding tissue, as demonstrated with a picosecond infrared laser (PIRL) (Franjic & Miller, 2010; Cowan *et al*, 2005; Böttcher *et al*, 2013, 2014, 2015). Previous studies have demonstrated the fundamental advantages of cutting tissue with a PIRL at 2940 nm wavelength compared to conventional CO_2_ lasers (Böttcher *et al*, 2014). These include almost scarless cutting, reduced thermal injury and high number of intact biomolecules in the arising aerosol, making it highly suitable for subsequent mass spectrometric analyses (Kwiatkowski *et al*, 2015; Wurlitzer *et al*, 2020; Hahn *et al*, 2021). Therefore, tissue sampling by IRL for proteomic or lipidomic analysis is a promising method, because it is both rapid and accurate and has been successfully used in various experiments (Saudemont *et al*, 2018; Ogrinc *et al*, 2019; Woolman *et al*, 2020a; Stadlhofer *et al*, 2023; Woolman *et al*, 2019; Voß *et al*, 2022; Hahn *et al*, 2021). Another advantage of this approach is the low sample volume, which allows precise and targeted sampling, enhancing the suitability for clinical use. For integration into clinical workflows, the laser irradiation should ideally be delivered via a handheld applicator, commonly achieved using a long and thin laser fiber, that can easily be installed in application-specific designs (Woolman *et al*, 2017a; Saudemont *et al*, 2018; Woolman *et al*, 2020b; Ogrinc *et al*, 2021, 2022; Woolman *et al*, 2024). Using this approach different types of pediatric brain cancer could be rapidly distinguished by a handheld PIRL coupled with real-time MS, termed PIRL-MS (Woolman *et al*, 2019, 2020b, 2024). “SpiderMass” represents another promising, minimally invasive technique that uses ambient MS for real-time analysis. This approach utilizes a nanosecond IRL (NIRL) coupled to a laser fiber, leveraging the principle of water-assisted laser desorption/ionization (WALDI). Using this method, Ogrinc et al. successfully differentiated non-tumorous, dysplastic, and tumorous areas of oral tongue squamous cell carcinoma (OTSCC) (Saudemont *et al*, 2018; Ogrinc *et al*, 2022).

OTSCC is a subtype of head and neck squamous cell carcinoma (HNSCC), which represents a heterogeneous group of malignant neoplasms, arising in the oral cavity, pharynx, and larynx (Johnson *et al*, 2020). Although the cancer originates from the squamous epithelium in all cases, oropharyngeal squamous cell carcinoma (OPSCC) in particular differs in terms of epidemiology, formation, risk factors and prognosis (Johnson *et al*, 2020; Ferris & Westra, 2023; Siegel *et al*, 2024). The incidence of OPSCC increased by 2.3% annually between 2009 and 2019 (Marur *et al*, 2010; Chaturvedi, 2012; Siegel *et al*, 2024). This trend is mainly attributed to increased infections with oncogenic human papillomavirus (HPV) type 16, which is present in more than 70% of OPSCC cases (Johnson *et al*, 2020). The lipidome of HNSCC has already been studied by different research groups, who identified specific increased or decreased lipid classes or species, that appear to be characteristic for tumorous and non-tumorous tissue (Schmidt *et al*, 2020; Yu & Wang, 2021; Dickinson *et al*, 2020; Stadlhofer *et al*, 2023). Although the analysis of OPSCC’s lipidome has been rarely performed, our recent study contributed to this emerging field by analyzing the lipidome of OPSCC in four HPV positive patients compared to each patient’s adjacent healthy tissue, using a NIRL-MS platform (Stadlhofer *et al*, 2023).

Building on these findings, we further expand this research in the present study using a larger sample set exclusively derived from the palatine tonsil, to minimize interpatient differences, using the established NIRL-MS platform. Both, HPV-positive and HPV-negative samples, were included to provide deeper insights into the differing pathogenesis and prognosis associated with HPV status. As a step towards *in vivo* tissue sampling, we demonstrate the application of a custom-made handheld laser fiber approach, which offers a more flexible and adaptable design compared to the conventional approach. Here, our aim is to identify potential lipid markers in low-volume OPSCC samples using both sampling approaches and to establish a reference databank, which is essential for online MS platforms such as SpiderMass and a future PIRL-MS platform.

## Results

In total, 28 samples from 11 patients (Table 1) were collected from the biobank. For each patient, one to three OPSCC samples and one adjacent healthy tissue sample (OPSCC n=17; healthy tonsil tissue n=11) were obtained. We included eight HPV-positive patients (HPV+ OPSCC n=12; healthy tonsil tissue n=8) and 3 HPV-negative patients (HPV-OPSCC n=5; healthy tonsil tissue n=3) in our study (Table 2).

**Table 1.** Demographic and clinical data of all patients. p16: p16 INK4A immunohistochemistry; HPV: human papilloma virus polymerase chain reaction for virus serotype 16 or 35.

|  |  |
| --- | --- |
| Number of patients (n) | 11 |
| Sex (n (%)) |  |
| male | 8 (72.7%) |
| female | 3 (27.3%) |
| Mean age at surgery (years) | 64.55 |
| Age range (years) | 47 - 79 |
| p16/HPV positive | 8 (72.7%) |
| p16/HPV negative | 3 (27.3%) |

**Table 2.**
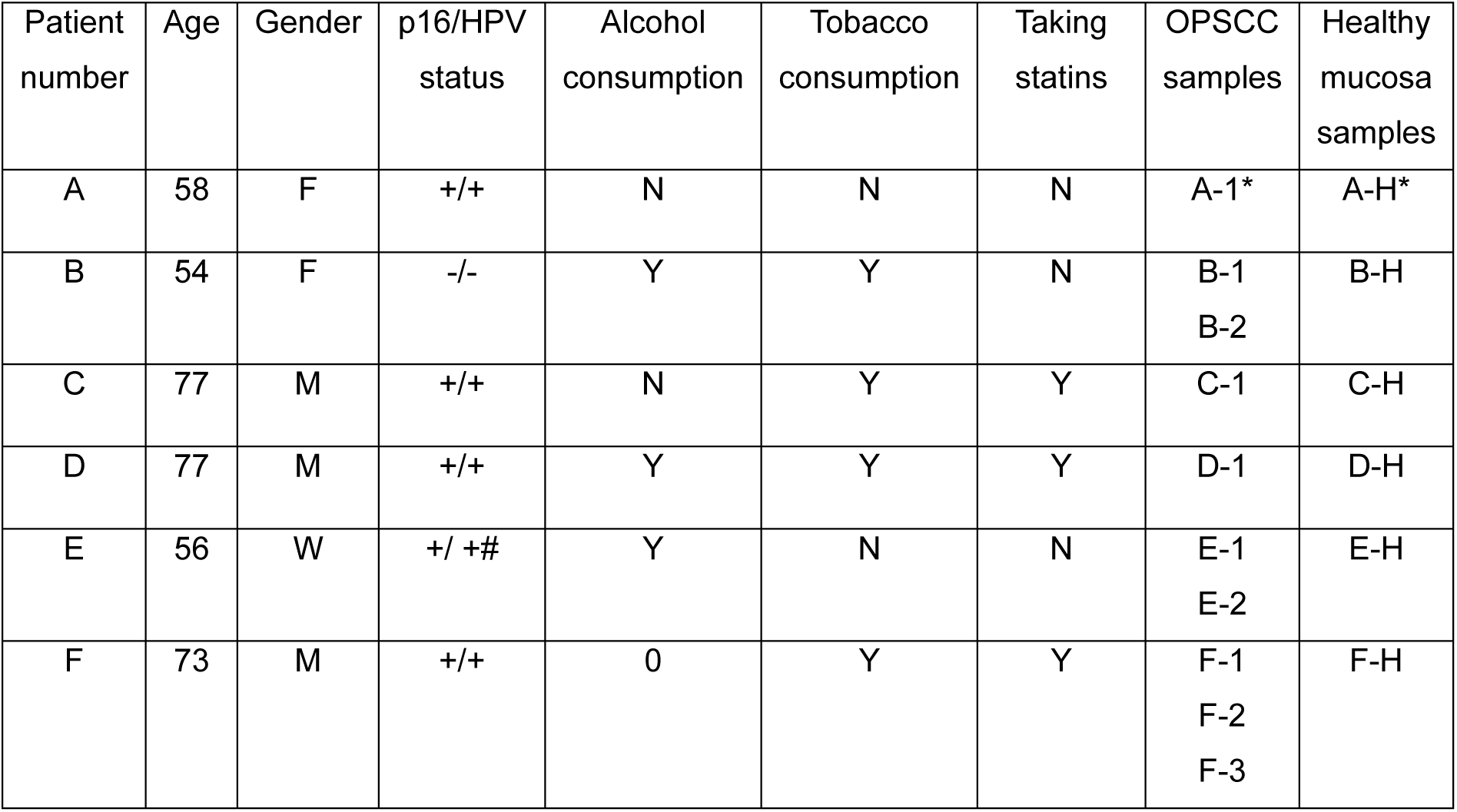
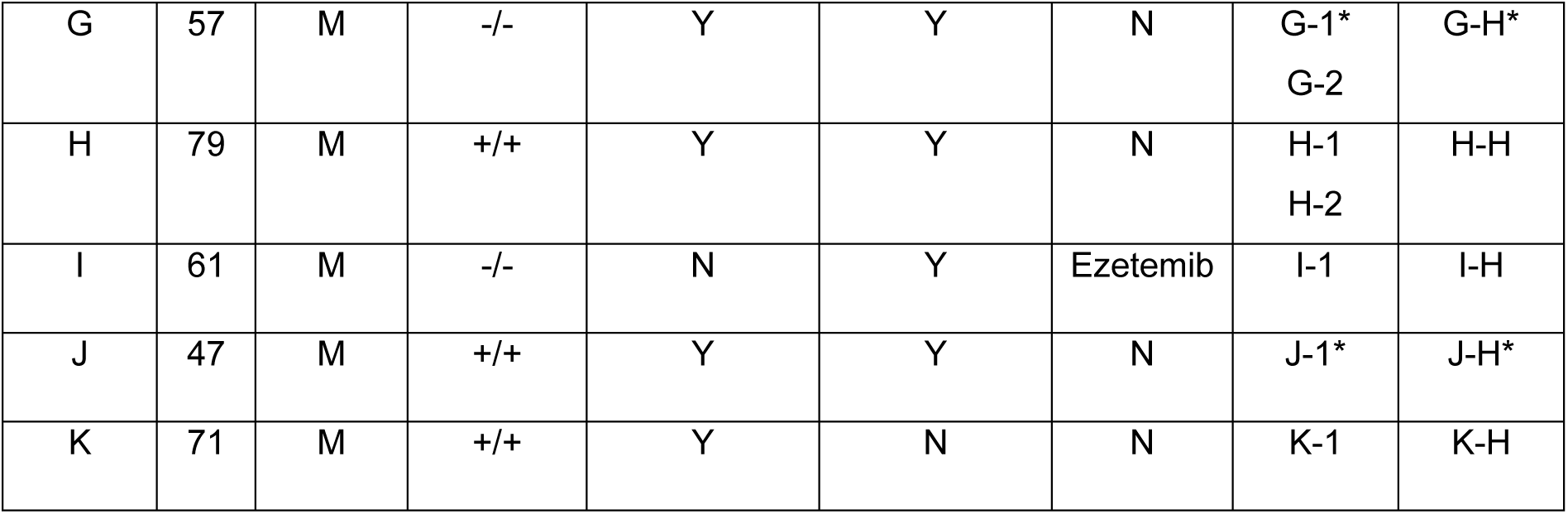
Detailed demographic and clinical data of all patients. p16: p16 INK4A immunohistochemistry; HPV: human papilloma virus polymerase chain reaction for virus serotype 16; # human papilloma virus polymerase chain reaction for virus serotype 35; *: samples that were ablated a second time with the handheld applicator.

The complete workflow of the study is depicted in Figure 1. The samples were irradiated using NIRL in two setups: (a) the chamber setup and (b) the handheld setup. The resulting tissue aerosol, containing intact biomolecules, was collected on glass fiber filters and further processed with the SIMPLEX protocol to extract lipids for shotgun lipidomics. Subsequently, data analysis was performed on the quantified lipid species concentrations using Welch’s t-test and orthogonal partial least square discriminant analysis (oPLS-DA) to identify lipidome differences and potential lipid biomarkers for future reference.

**Figure 1.**
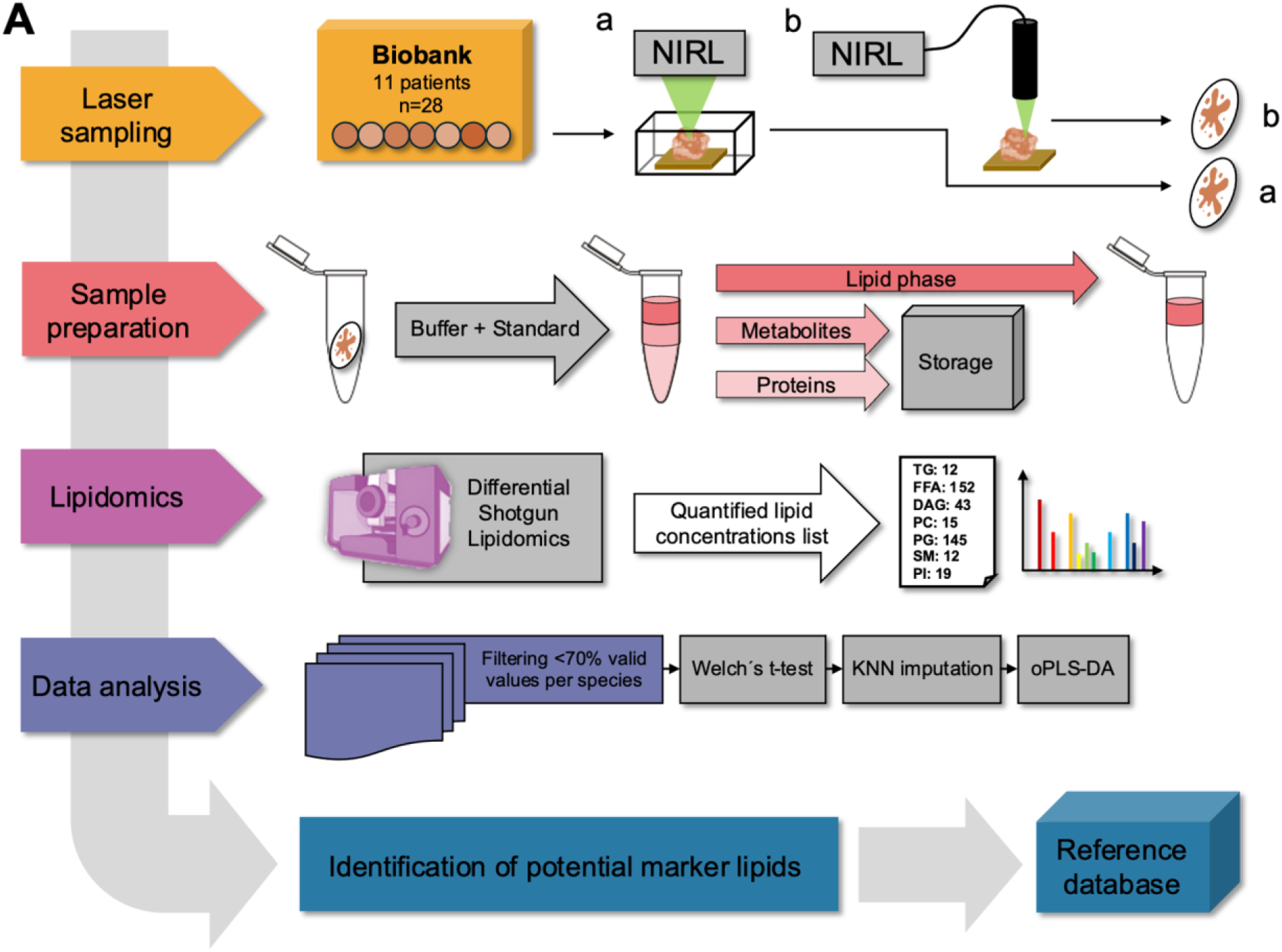
**A** Experimental design and data analysis scheme of the study

Each sample was ablated three times (technical triplicates) in the closed chamber of the stationary ablation setup, resulting in a total of 84 mass spectrometric measurements. In addition, six samples from three different patients (OPSCC n=3; healthy tonsil tissue n=3) were ablated a second time (technical duplicates) using a handheld applicator developed in-house for comparison with the chamber MS results. Technical triplicates were prepared in the same way.

### Distribution of all lipid classes from the chamber setup

To gain an overview of the OPSCC and tonsil lipidome across the patient cohort prior to detailed analysis, the absolute lipid class concentration and lipid class composition of each individual patient were analyzed. In total, we identified 885 lipid species across 16 lipid classes, including: Cholesterol Ester (CE); Ceramides (Cer d18:0, Cerd18:1); Diacylglycerides (DG); Free Fatty Acids (FFA); hexosylceramides (HexCER); lysophosphatidylcholine (LPC); lysophosphatidylethanolamine (LPE); Lactosylceramide (LacCER); Phosphatidic Acid (PA); Phosphatidylcholine (PC); phosphatidylethanolamine (PE); Phosphatidylglycerol (PG); Phosphatidylinositol (PI); Phosphatidylserine (PS); Sphingomyelin (SM); and triacyltriglycerides (TG).

The relative proportion and the log_2_-transformed lipid concentrations of representative Patients A, B, and F are shown in Figure 2. All other patients’ individual lipid class concentrations are provided in Supplementary Tables S1a and S1d. Patient A and F have positive HPV-status and Patient B has negative HPV-status. In Patient A, the lipid classes CER, LPE, LacCER, PC and PG are clearly elevated in the OPSCC tissue, with LacCER showing the highest relative increase. The most altered lipid class is TG, which appears at markedly lower concentrations in the OPSCC sample. The difference is particulary noticeable, as TG accounts for 4.43% in sample A-1, but almost 85% in the adjacent healthy tissue (sample A-H). The combined proportion of PC and PG decreases notably from 35% to under 5% in the healthy tissue. All other lipid classes are either slightly increased or remain unchanged in their concentration.

**Figure 2.**
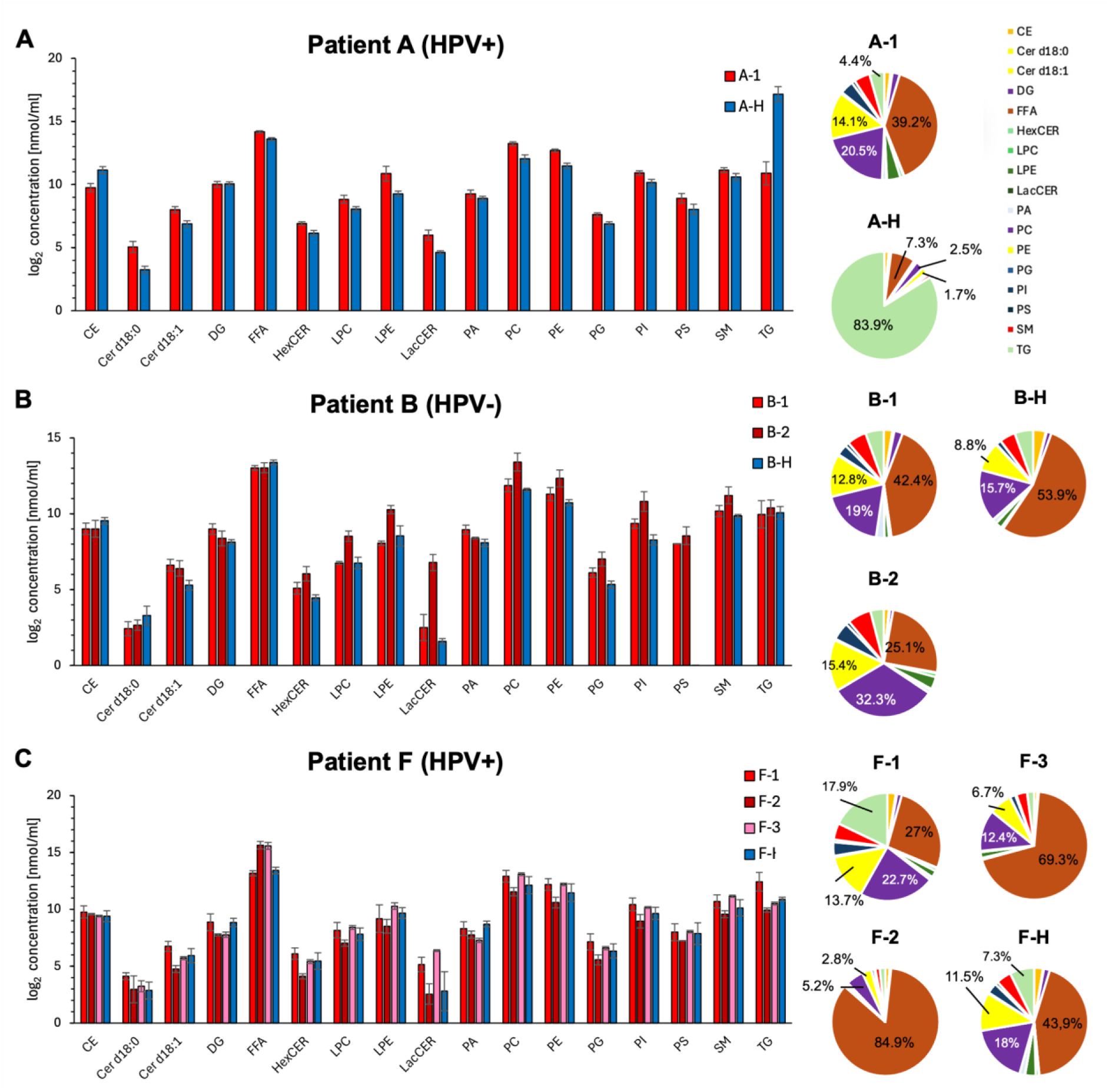
**A** Individual log_2_ lipid class concentration and relative proportion of lipid classes of Patient A. The error bars show the standard deviation of the lipid concentration of each patient calculated by the three technical replicates. **B** individual log_2_ lipid class concentration and relative proportion of lipid classes for Patient B. **C** Individual log_2_ lipid class concentration and relative proportion of lipid classes for Patient F.

In Patient B, FFA constitute the largest lipid proportion in both, OPSCC (B-1) and healthy tissue (B-H), while PC dominates in sample B-2. The lowest proportions of PC and PE are seen in the healthy samples, mirroring the pattern of Patient A, though less distinctly. In contrast to Patient A, some lipid classes are decreased in the OPSCC samples of Patient B, including CE, Cer d18:0 and FFA. TG remains unchanged in its concentration. However, LacCER is strongly increased, especially in sample B-2, similar to the trend in Patient A. Additional increases are seen for HexCER, PC, PG and PI, all of which are more prominent in sample B-2.

Across all patients, a general trend toward increased lipid class concentrations in OPSCC was observed. Notably, LacCER consistently increased across all patients, except in Patients F and K. HexCER, LPE, PC, PE and PG also showed higher median concentration in OPSCC tissue in most patients, with the exception of Patients C and F. Furthermore, Cer d18:1 was increased in all patients, except Patient D. In contrast, TG was the only lipid class exhibiting a clear decrease in multiple OPSCC samples, specifically reduced in Patients A, E, I, G, J and K.

As contrasting results were observed in Patient F, we examined its lipid distribution in more detail. Patient F had three OPSCC samples, with samples F-1 and F-H taken during panendoscopy and samples F-2 and F-3 collected three weeks later during tumor resection. Samples F-2 and F-3 showed unusually high proportions of FFA (84.88% in F-2 and 69.38% in F-3). A general pattern of lipid alteration was difficult to discern in Patient F. Some lipid classes showed large differences, but these were often limited to individual samples. For example, TG was increased only in sample F-1, while HexCER was decreased only in sample F-2. LacCER was increased in sample F-1 and F-3. The similarity between samples F-1 and F-H was greater than between OPSCC samples F-1, F-2 and F-3, possibly due to temporal lipidome fluctutations or differences in sample handling. As the high amount of FFA >80% in F-2 suggests the occurrence of lipolysis, this sample is excluded from further analysis.

For an overview of lipid class distribution across the whole patient population, the distribution of six lipid classes with a high difference in median concentration between healthy and OPSCC samples is visualized as Violin Plots in Figure 3, revealing concentration differences between the tissue types in these lipid classes.

**Figure 3.**
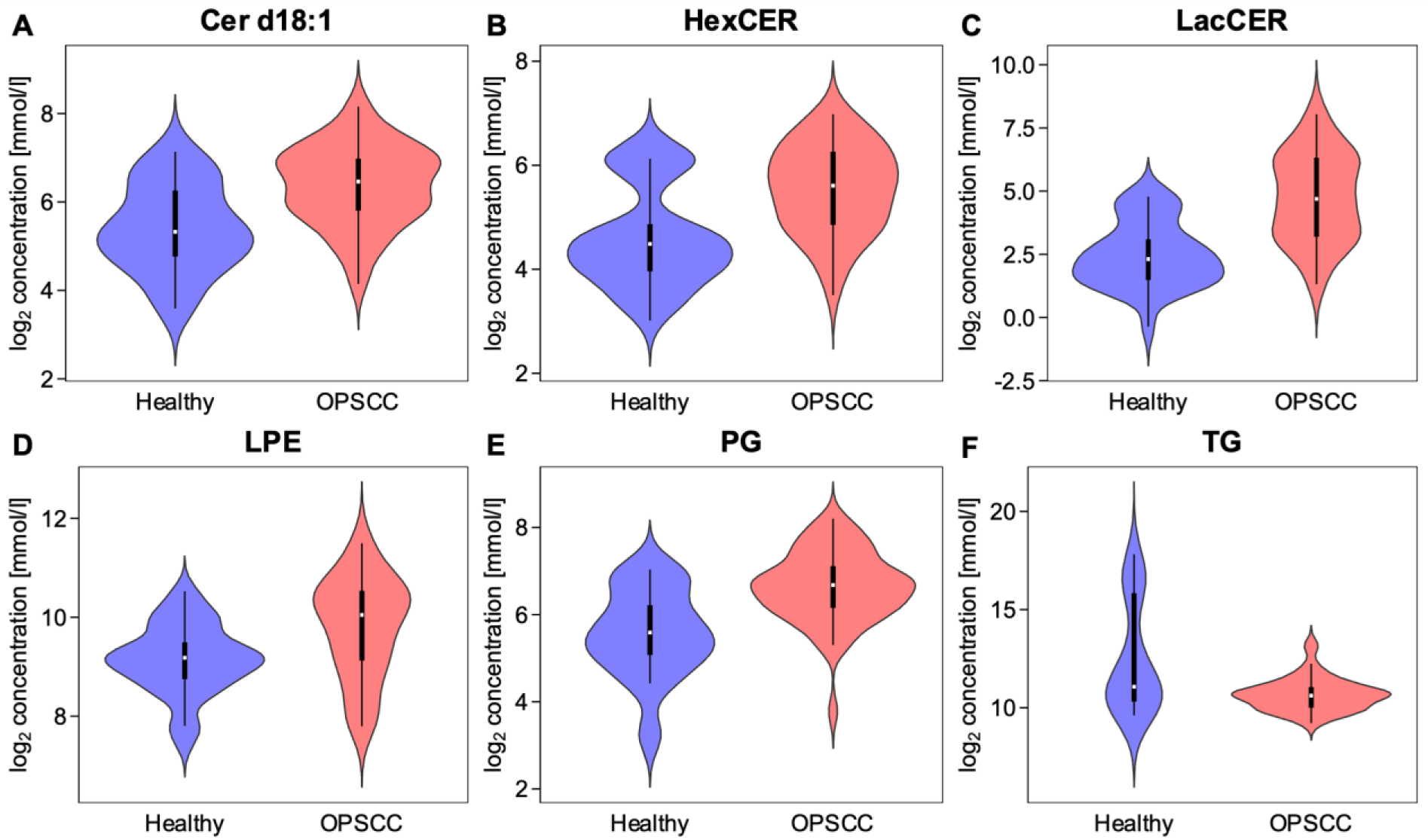
**A - F** Violin plots show the log_2_ concentration of lipid classes with high difference in median concentration across all healthy and all OPSCC samples. Box plots within the violin plots representing the interquartile range (first quartile (Q1) to third quartile (Q3)), containing the middle 50% of the data. The white line inside the box indicates the median. Extend lines are drawn to the maximum and minimum measured concentration excluding outliers lower or greater than Q1 - 1.5 * IQR or Q3 + 1.5 * IQR.

Higher median concentration in the OPSCC samples can be seen especially in the lipid classes Cer d18:1, HexCER, LacCER, LPE and PG. Median concentration of LacCER is more than fourfold higher in OPSCC (log_2_ concentration difference: 2,086 mmol/l), with the highest difference across all lipid classes. Nevertheless, this lipid class shows a very high variance across all samples, with a high interquartile range (IQR) and large difference between the minimum and maximum measured concentration in both, OPSCC samples and healthy samples. Cer d18:1, HexCER and PG have the next highest median differences (log_2_ concentration difference: >1 mmol/l). Their IQRs overlap between the entities and maximum and minimum concentrations span a broad range. LPE has a nearly completely overlapping IQR, but also relatively high difference in median (log_2_ concentration difference: 0.83 mmol/l) compared to the other lipid classes. TG is the only lipid class, that has a clearly lower median concentration (log_2_ concentration difference: −0.334 mmol/l) in the healthy samples. Noteworthy is that TG have a low IQR in the OPSCC samples, while showing a very high IQR in the healthy samples. The first quartile (Q1) and minimum concentrations of both entities are in a similar range. Meanwhile, the third quartile (Q3) of healthy samples is higher than the maximum detected concentration in OPSCC samples. The maximum concentration is approximately eightfold higher in healthy samples than in OPSCC samples, indicating that TG concentration varies more in healthy tissue than in OPSCC tissue, with outliers exhibiting high concentration in healthy tissue.

In general, most lipid classes show small distance in concentration between the first quartile and the third quartile. The low IQR indicates that the lipid class is often concentrated similarly within one entity. Lipid classes with very low IQR are CE, DG, FFA, LPC, PA, PS and SM (log_2_ IQR <1,2mmol/l). All lipid classes except TG and FFA tend to have a higher median concentration in OPSCC. We found no lipid class, that allowed a clear distinction across all patients.

### Differentiation between healthy and tumorous tissue

To evaluate the particular lipid species differences between healthy and tumorous tissue, Welch’s t-test (p-value < 0.05, two-fold change) was performed for each individual patient on data filtered for 70% valid values. Missing values were imputed using the [k-nearestneighbors](https://www.geeksforgeeks.org/k-nearest-neighbours/) (kNN) algorithm (k=5). In total, 532 lipid species were included in the analysis. The results are provided as table in Supplementary 4. The volcano plots for the selected representative Patients A, B, F and K are presented in Figure 4. Patient A, with HPV positive status has 328 (281 decreased, 47 increased) significantly altered lipid species in sample A-1. Sample A-1 contains a large number of significantly decreased TG species, but lipid species from the classes CE, DG, HexCER, PG and PI are also significantly decreased in this patient. The increased lipid species are distributed across several lipid classes, and almost every lipid class has at least one significantly altered lipid species in this patient. In Patient K, a very similar pattern is observed. The significantly decreased lipid species are again dominated by TGs, while the increased lipid species show a diverse composition. We found this pattern in 7 of 11 patients in at least one sample (Patient A, E, G, H, I, J, K) with varying number of decreased and increased lipid species.

**Figure 4.**
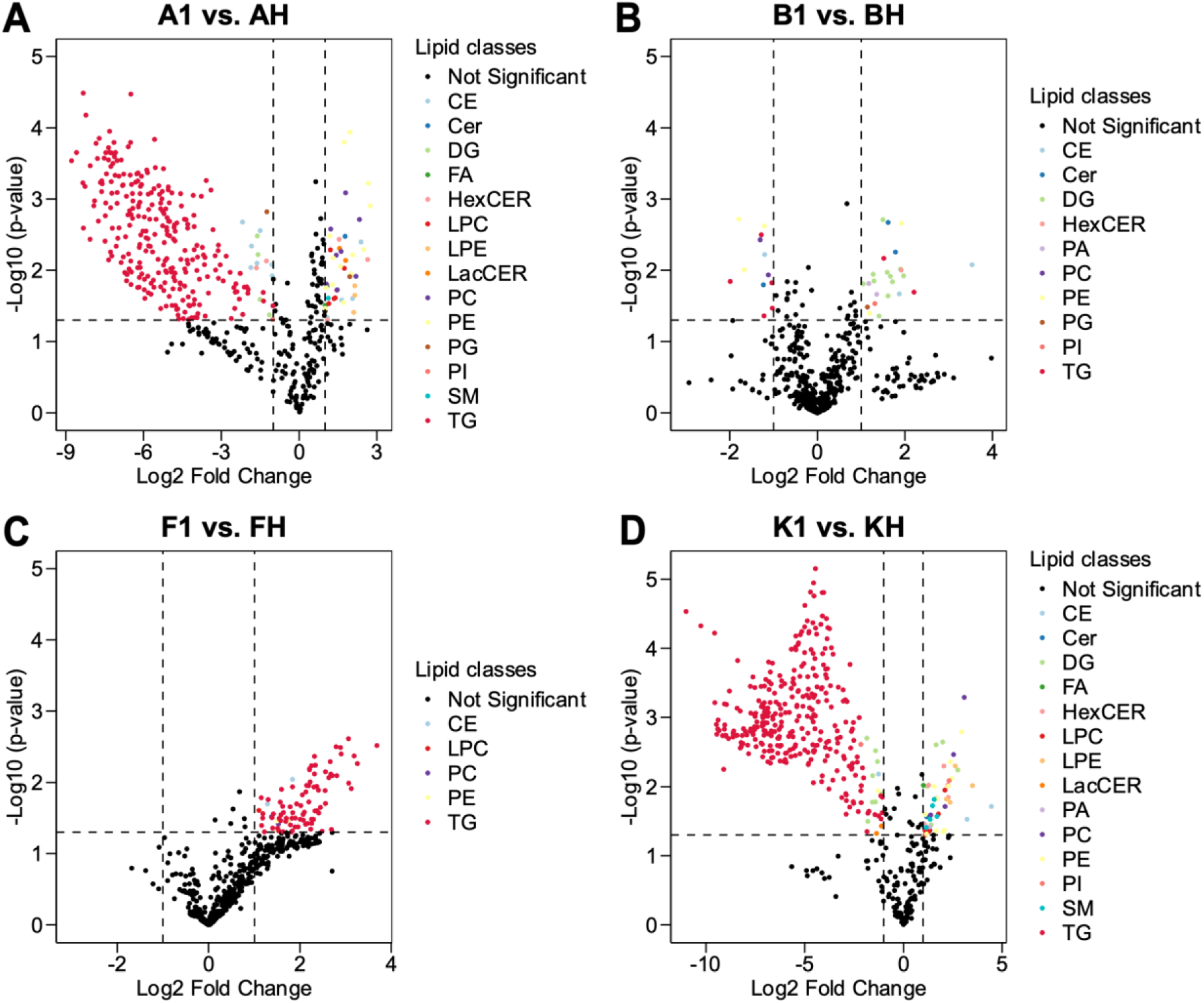
**A** Results of Welch’s t-test visualized as Volcano Plot calculated from three technical replicates from sample A-1 versus sample A-H. Significantly increased or decreased lipid species are highlighted according to their lipid class (p-value < 0.05, two-fold change). Lipid species, which are not significantly altered (p-value < 0.05 and/or log_2_ fold change <1) are colored black. **B** Volcano plot from sample B-1 versus B-H. **C** Volcano plot calculated from sample F-1 versus F-H. **D** Volcano plot calculated from sample K-1 versus K-H.

Patient B, who has HPV negative status, has 12 decreased lipid species and 24 increased lipid species in sample B-1. CE, CER, PC, PE and TGs have significantly decreased lipid species, while several different lipid species from nine different classes are significantly increased. Sample F-1 has only significantly increased lipid species (91 increased) compared to sample F-H. In contrast to the other patients, these are mainly TG species with high log_2_ fold change.

### Lipidome analysis of handheld-ablated tissue samples

In total, six samples from three different patients were ablated a second time using the new handheld applicator. With the handheld experimental setup, 16 different lipid classes consisting of 858 lipid species were detected. All lipid classes that were identified in the conventional chamber setup were also detected using the new handheld applicator. The total number of detected lipid species was only slightly lower with 885 lipid species identified in the conventional setup. Overall, the quantified concentrations were lower when using the handheld applicator.

The individual lipid class concentrations of the three patients are shown in Figure 6. In the OPSCC samples from Patient A, most lipid classes are present in higher concentrations, only the TGs are present in slightly lower concentrations. Notable differences are observed in the concentrations of Cer d18:1, DG, FFA, HexCER, LPC, LPE, LacCER, PC, PE, PG and PI. The decrease in TG levels in Patient A is minimal. Consistent with the chamber ablation results for the samples from Patient A, PC and PG show a reduced proportion in the healthy tissue. The variation within lipid class concentrations is low, as indicated by a low standard deviation. In Patient G, the lipid classes CE, Cer d18:1, HexCER, LacCER, PA, PC, PG, PI, PS and SM are increased in the OPSCC samples. A particularly high relative change is observed for TG. Also, the concentration of TGs in the healthy sample G-H is the highest of a lipid class in the handheld experimental setup. DG is decreased in the OPSC sample of this patient. Again, PC and PE exhibit reduced proportions in the healthy sample. In contrast to patients A and G, no lipid class is either clearly increased or decreased in Patient J. Although TG species have the highest relative change, the technical replicates of the healthy sample show high variability, which precludes a definitive interpretation.

**Figure 5.**
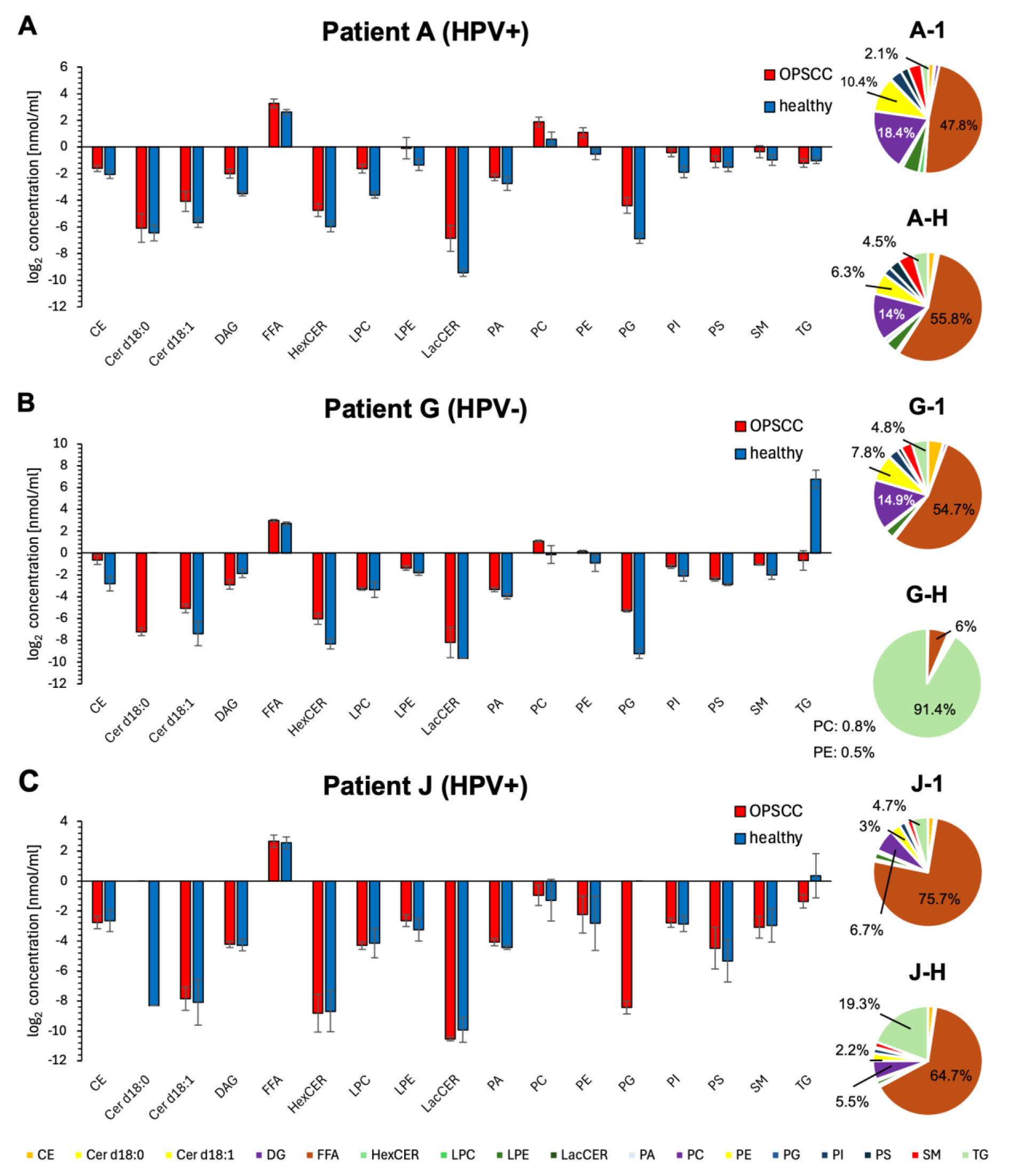
**A** Individual log_2_ lipid class concentration and relative proportion of lipid classes sampled with the handheld applicator from Patient A. The error bars show the standard deviation of the lipid concentration of each patient calculated by the three technical replicates. **B** Individual log_2_ lipid class concentration and relative proportion of lipid classes from Patient G. **C** Individual log_2_ lipid class concentration and relative proportion of lipid classes from Patient J.

**Figure 6.**
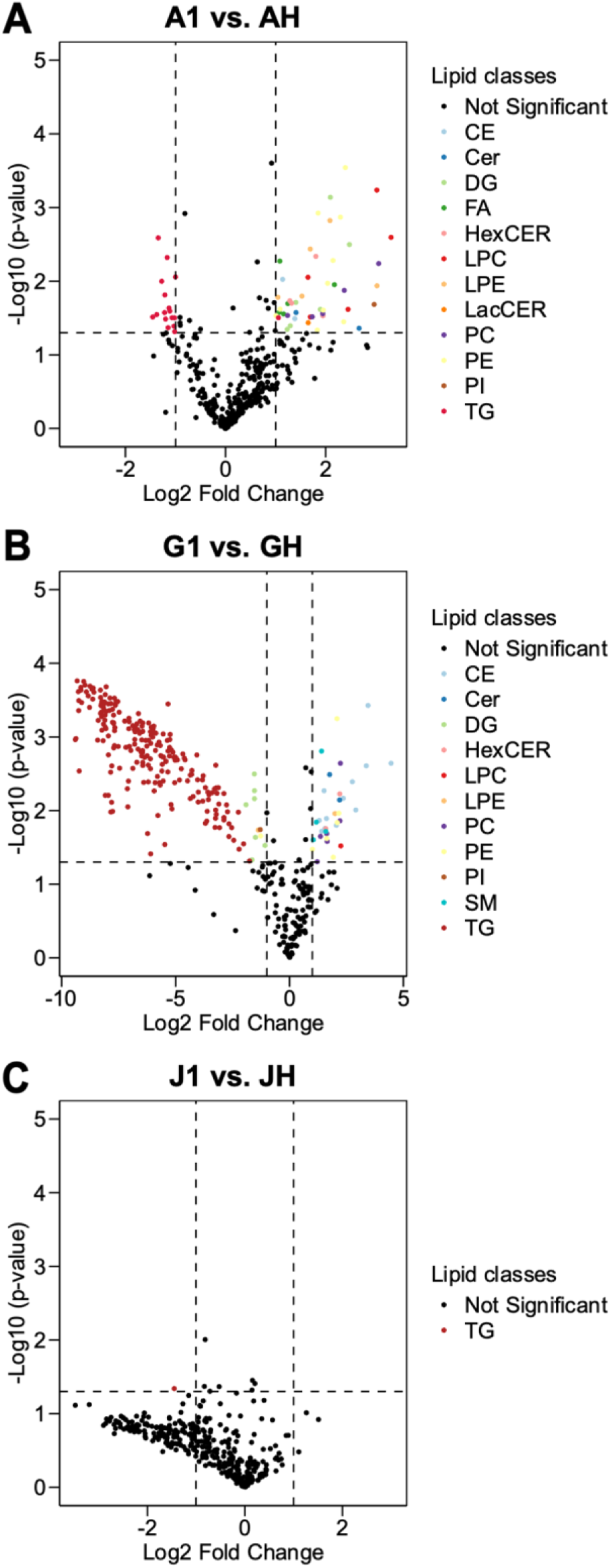
**A** Results of Welch’s t-test visualized as Volcano Plot from three technical replicates from sample A-1 versus sample A-H sampled by the handheld applicator. Significantly increased or decreased lipid species are highligted according to their lipid class (p-value < 0.05, two-fold change). Lipid species that are not significant altered (p-value < 0.05 and/or log_2_ fold change <1) are colored black. **B** Volcano Plot from sample G-1 versus G-H. **C** Volcano Plot from sample J-1 versus J-H.

### Differentiation between healthy and tumorous tissue with handheld applicator

After filtering for 70% valid values and performing kNN imputation (k = 5), Welch’s t-test (p-value < 0.05, two-fold change) was performed including 401 lipid species for the three patients individually; the results were visualized as volcano plot (Figure 6). In Patient A, 16 lipid species were significantly decreased and 47 significantly increased. The decreased lipid species were exclusively TG species. The increased lipid species are made up of 12 different lipid classes, with the highest number of PE lipid species. Patient G exhibited 266 (232 decreased, 34 increased) significantly altered lipid species, which is the highest number of the three patients. Among the 232 significantly decreased lipid species, 222 belong to TG, while the remaining were predominantly DG species. The increased lipid species are composed of eight different lipid classes, most of which are either CE, PC or PE. Of particular note is the high log_2_ fold change observed in many TG species, reaching values up to −10. In Patient J, only TG 54:5-FA22:5 was significantly decreased; all other lipid species showed no significant alterations in its concentration. Nevertheless, Patient J also showed decreased concentrations of TGs with a high log_2_ fold change, though these were not statistically significant due to high variability among the technical replicates.

Commonalities in lipid profiles were observed across the patients, especially between Patient A and G, with ten identical lipid species significantly increased and 16 identical lipid species significantly decreased in the OPSCC sample. The significantly decreased TG 54:5-FA22:5 in Patient J was also found to be decreased in Patient G.

### Identification of potential OPSCC marker lipids from the chamber setup

We performed orthogonal partial least square discriminant analysis (oPLS-DA) including 532 lipid species (filtered for 70% valid values; kNN imputation (k=5)) to identify the lipid species contributing the most to differentiation between OPSCC and healthy tissue. Technical replicates were averaged. The scatter plot in Figure 7 shows a clear separation of healthy tissue and OPSCC tissue by the model along the principal component. The OPSCC samples show a distinct grouping while the healthy samples are more broadly distributed along the orthogonal component. The 95% confidence intervals are not overlapping, indicating that the model differentiates well between both groups. The lipid species contributing the most to tissue differentiation are mainly CE, HexCER and PE. All lipid species from CE, HexCER, LPE, PC, PE and PG are present exclusively at increased concentrations in the OPSCC tissue, indicating that increased lipid species concentrations are the most important hallmark for OPSCC. Model validation using 100 permutations yielded Q^2^=0.753 with a permutation p-value < 0.01 and R^2^Y=0.887 also with a permutation p-value < 0.01. The robust R2Y and Q2 values, confirmed through permutation testing, demonstrate the model’s effectiveness in capturing the variance within the dataset and its strong predictive power. The significant p-values further support the reliability of the results. The highlighted lipid species have a high potential to be future markers for OPSCC tissue differentiation and warrant dedicated focus in further analysis to fully explore their diagnostic potential.

**Figure 7.**
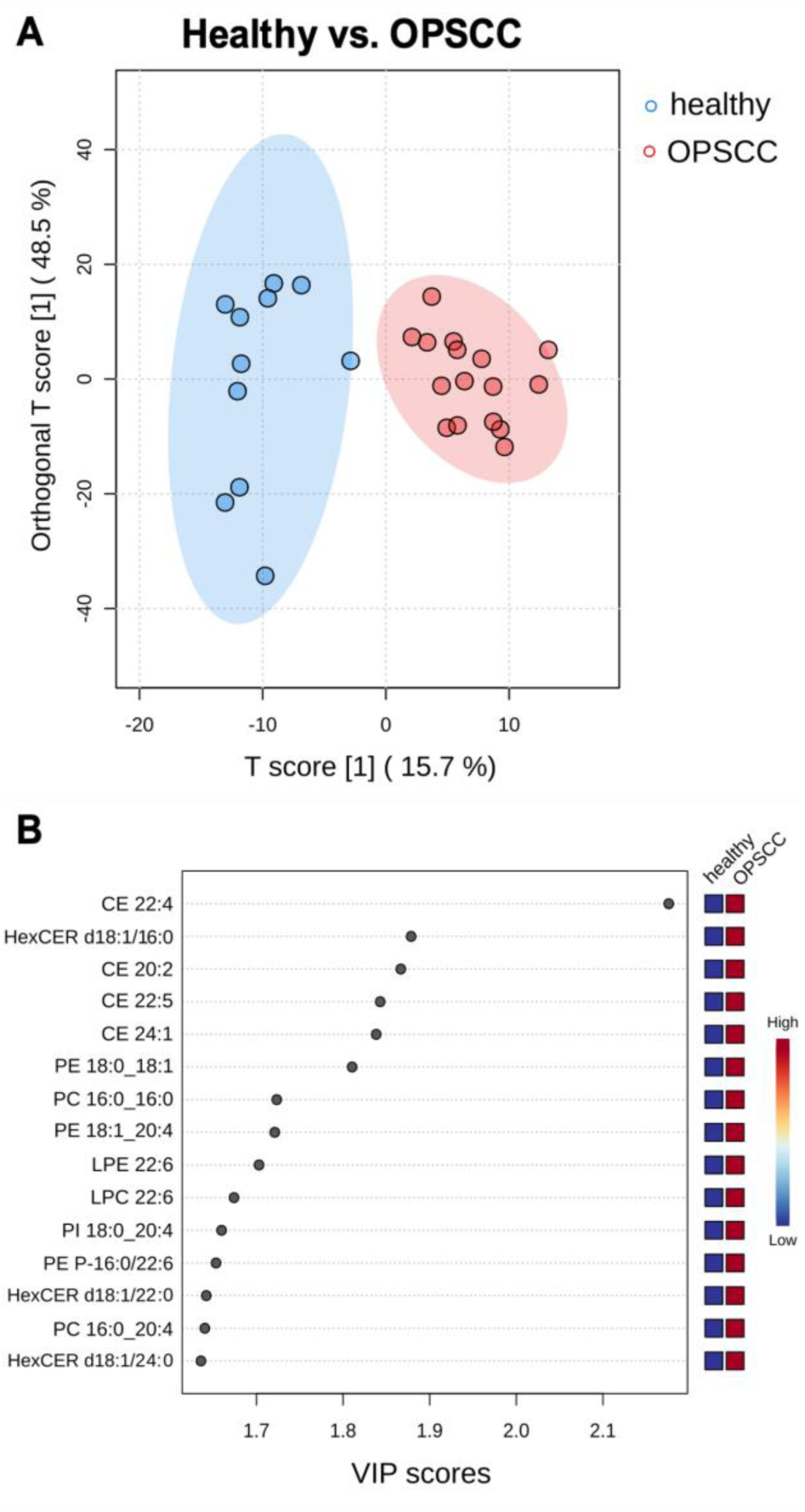
**A** Results of oPLS-DA visualized as scatter plot. Technical replicates were averaged before performing oPLS-DA. Each dot represents one sample. **B** VIP score plot of lipid species sorted after importance for tissue differentiation.

For further validation and to identify additional lipid species that may serve as potential OPSCC markers, the results of Welch’s t-test were analyzed collectively across all samples, as illustrated in Figure 8. A summary of this analysis is provided in Supplementary 4a. Twenty lipid species were identified, showing significant increases or decreases in at least 8 and up to 13 out of 16 samples within the chamber setup. CE 22:4, CE 22:5, HexCER d18:1/22:0 and PE 18:0_18:1 were the most frequently significantly altered lipid species, increased in 11, 12 or 13 of 16 samples. At least one of these species is significantly altered in every sample analyzed within the chamber setup. These species were among the top 20 most important lipid species in oPLS-DA, too. As shown in Figure 8, they were also detected by the handheld applicator and similarly altered as in the chamber study. Although CE 22:4 was excluded from performing Welch’s t-test (<70% valid values) in the handheld study, it was still detected using the handheld applicator almost exclusively in the OPSCC samples, making it a promising lipid species for differentiation. In Figure 8C-H, Box Plots visualize the distribution of the three most promising lipids separated by chamber and handheld study. All plots show a clear differing concentration between healthy and OPSCC samples. The behavior of these lipid species corresponds well between both studies, proving the function and future potential of a handheld applicator.

**Figure 8.**
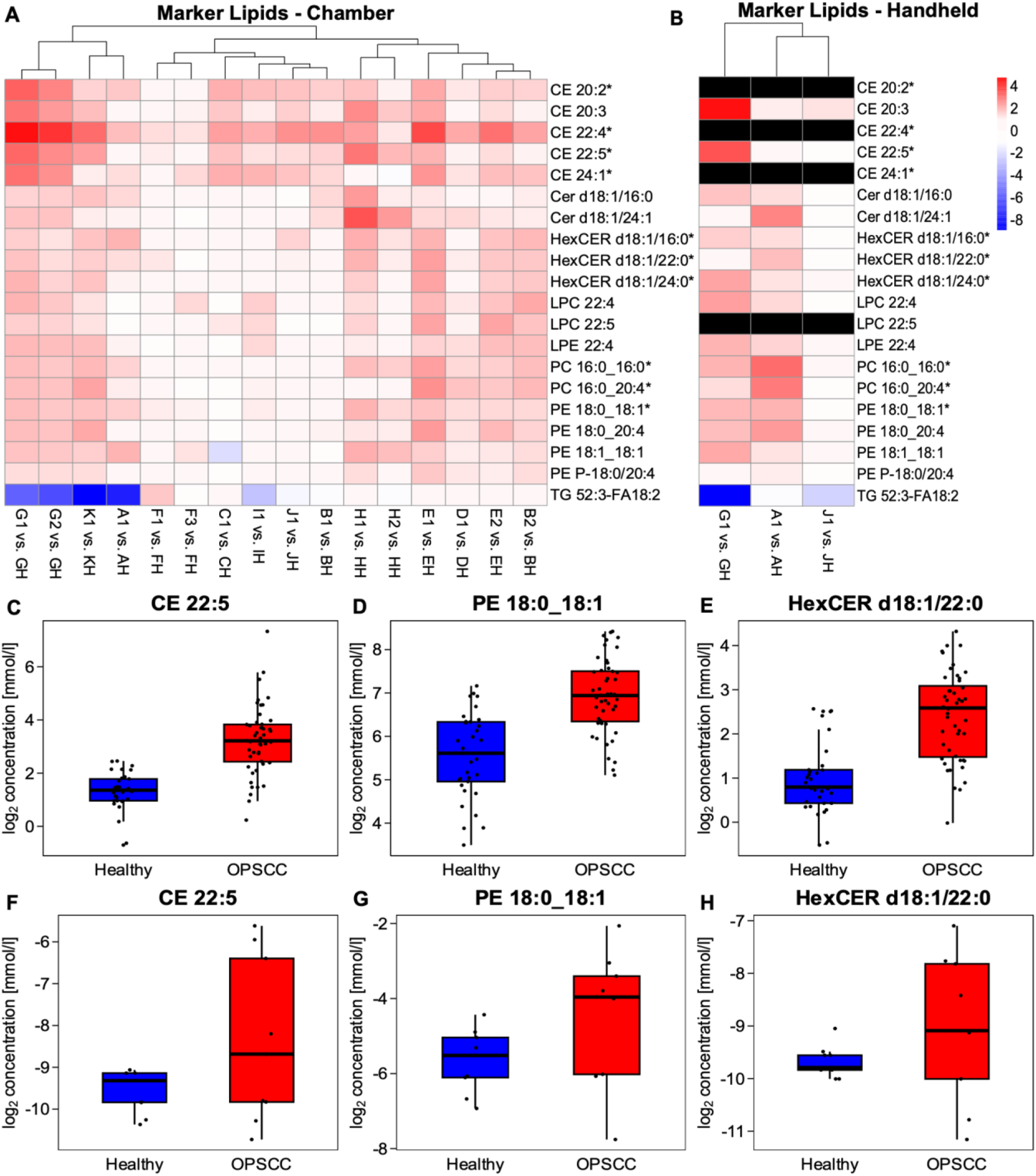
**A** log_2_ fold change (healthy vs. OPSCC) of 19 most often significantly (p=0.05, two-fold change) increased lipid species and one representive TG in the chamber setup visualized as heatmap. Species having a high VIP score in oPLS-DA (Top 15) are marked with *. **B** log_2_ fold change (healthy vs. OPSCC) of these lipid species, ablated by the handheld applicator. Species detected in under 70% of the samples are excluded (black) **C D E** Box Plot of one lipid species per class ablated by the chamber setup, that fulfill following criteria: significantly increased in more than 10 samples, Top 15 in oPLS-DA and incldued in the handheld analyis **F G H** Box Plot of the species which met the criteria sampled by the handheld applicator.

## Discussion

The results of our study, following the approach of rapid tissue identification, demonstrate the feasibility to distinguish OPSCC from healthy tissue to enhance the precision of surgical resection. Our analysis revealed similar OPSCC lipid profiles in a subset of patients and identified significant alterations in several lipid species, which, in combination, are consistently present across all samples, independent from age, sex, HPV status and smoking behavior, highlighting their potential as marker lipids. Our findings have been validated by both, oPLS-DA and Welch’s t-test, providing robust evidence to support the potential of these lipids as future biomarkers. Aiming to expand on previous research, we included a larger and more diverse patient population, incorporating both, HPV-positive and HPV-negative samples. Using an established chamber setup, we conducted lipidomic analysis on low volumes of 180 nL of fresh frozen tissue samples. We refined our sampling method to obtain a more representative overview by ablating four different locations per replicate. With this study, a lipidomic database is further established for healthy and OPSCC tissue of the palatine tonsil and marker lipids for OPSCC tissue are identified, that could be applied in both scientific research and diagnostic settings in the future. Furthermore, we achieved comparable results using a custom-made handheld applicator, representing an advancement in our sampling method and demonstrating its potential for more flexible and practical applications.

Lipidomics alterations have been documented across various cancer types, including lung, breast, gastrointestinal and prostate cancer, among others (Hung *et al*, 2019; Li *et al*, 2016; Ecker *et al*, 2021; Cífková *et al*, 2015; Eggers *et al*, 2017). Lipid changes vary considerably between cancer types. For example, in contrast to our findings, increased TG concentrations have been reported in colorectal cancer (Ecker *et al*, 2021). Additionally, a study by Ogrinc et al. on oral tongue squamous cell carcinoma identified PC, PE, PA, PI and PS as characteristically altered lipid classes (Ogrinc *et al*, 2022). Similarly, in our study, PC and PE tended to be altered, with increased concentrations in OPSCC tissue. Consistent with our findings, Li et al. identified accumulated Cholesteryl Esters as characteristically for prostate cancer (Li *et al*, 2016). In our data, CE 22:4 and CE 22:5 emerged as highly characteristic lipid species for OPSCC. Additionally, PE 18:0_18:1, which was significantly increased in most OPSCC samples, was recently identified as early-stage marker lipid for lung adenocarcinoma (Sun *et al*, 2024). These findings demonstrate overlap with other studies, despite the use of diverse sampling and MS methods, while highlighting the novelty of our work as the first, to our knowledge, to confirm these lipid alterations specifically in palatine tonsil carcinoma.

Despite these promising results, our study has limitations that may affect the applicability of the findings. Inter- and intrapatient variability in the lipidome, already reported in other studies (Ogrinc *et al*, 2022; Stadlhofer *et al*, 2023), was also observed in our study. The consistency of the results was disrupted by samples that showed entirely opposite lipid changes, such as exclusively increased and decreased lipid species within a patient or only a single significantly altered lipid species in the handheld applicator experiment. We also observed temporal fluctuations in the lipidome in Patient F. One sample from the chamber study was excluded from further analysis, due to an abnormally high proportion of FFA, most likely resulting from lipolysis that occurred during sample handling. Nevertheless, the overall lipid patterns were generally consistent with our previous OPSCC study (Stadlhofer *et al*, 2023). Lipid profile deviations from the patterns measured in most patients can be explained by the heterogeneous composition of the samples. Macroscopically, it is impossible to definitively confirm whether the ablated tissue consists solely of tumor tissue or also includes healthy or dysplastic tissue. Due to the large number of samples analyzed, heterogeneity appears to be a hallmark of the lipidome of OPSCC, whether due to the tumor tissue itself, measurement errors or external patient-associated factors such as genetics, diet or smoking. To spatially resolve this heterogeneity, an adjacent healthy tissue sample or areal would be necessary for reference to detect the changes in the OPSCC tissue lipidome, which could be addressed in the future. Furthermore, the integration of additional omics data, such as proteomics and metabolomics, is feasible, as we successfully applied ‘Simultaneous Metabolite, Protein, Lipid Extraction’ (SIMPLEX) protocol in this study (Coman *et al*, 2016). This approach may enhance the tissue differentiation and provide deeper insights in the molecular composition of OPSCC.

Although the handheld applicator represents a step towards a more *in vivo* orientated sampling method, this approach requires some improvements beforehand. Laser ablation of the sample was performed on a computer-controlled movable cooling stage, allowing precise scanning according to predefined instructions. In the future, the surgeon or the applicator itself should execute the scanning movement to sample a representative tissue section. Additionally, the time required for ablation could be reduced by a factor of 50 to 75 by using a next-generation picosecond infrared laser (PIRL), offering a repetition rate of 1 to 1.5 kHz.

Our results suggest that an IRL-based handheld applicator could assist surgeons during cancer resection by sampling suspected tumor regions and margins, especially when combined with an online mass spectrometric lipid analysis. Nowadays the intraoperative assessment of resection margins is often accomplished by the surgeon based on subjective visual and tactile evaluation (Baddour *et al*, 2016). The objective confirmation is provided only by a pathologist, who examines fresh-frozen tissue sections (Baddour *et al*, 2016; Ribeiro *et al*, 2003). Although this is a highly validated method, it is time-consuming and may be compromised by disruptive factors that reduce the adequacy (Layfield *et al*, 2018; Black *et al*, 2006; Baddour *et al*, 2016). Therefore, it is desirable to simplify and accelerate this complex process, because a complete resection is a crucial factor for patient outcome (Baddour *et al*, 2016). Tools for margin evaluation, such as fluorescence-guided surgery that can stain the tumor during surgery, are a significant further development, that has advanced oncologic surgery (Stummer *et al*, 2006). In addition, ambient mass spectrometry could be such a tool that is not yet well established but has proven great potential (Fatou *et al*, 2016; Woolman *et al*, 2019; Balog *et al*, 2013; Chen *et al*, 2013).

In conclusion, we confirmed the feasibility of differentiating between fresh-frozen tissue samples from healthy and OPSCC tissue from the palatine tonsil in a larger patient population with different HPV statuses. Our study reveals distinct lipidome differences in OPSCC tissue and identifies potential characteristic OPSCC marker lipids, although these findings warrant further validation in larger independent cohorts. We further enhanced our conventional approach by developing a mobile and more realistic setup, demonstrating that tissue differentiation is also achievable using this method. Notably, the handheld applicator represents a significant step towards clinical application compared to the lab only chamber setup. Overall, our promising results lay a strong foundation for future clinical innovations, potentially adding a new pillar to intraoperative diagnostic in the future and contributing to the development of new diagnostic approaches in head and neck oncology.

## Materials and Methods

### Reagents and Tools Table

| Reagent/Resource | Reference or Source | Identifier or Catalog Number |
| --- | --- | --- |
| **Chemicals, Enzymes and other reagents** |  |  |
| *Ultimate SplashOne* | Sigma-Aldrich | 330820L-1EA |
| *oleic acid d9* | Sigma-Aldrich | 861809O-10MG |
| *Glu Cer(d18:1-d7/15:0)* | Sigma-Aldrich | 330726P-1MG |
| *dLacCER d18:1/15:0* | Sigma-Aldrich | 330729P-1MG |
| *15:0-18:1-d7-PA* | Sigma-Aldrich | 330727P-1MG |
| *dCer d18:0/13:0* | Sigma-Aldrich | 791642C-1MG |
| **Software** |  |  |
| *ITK-Snap* | (Yushkevich *et al*, 2006) | Version 3.8.0 |
| *Fiji (ImageJ)* | (Schindelin *et al*, 2012) | Version 1.54f |
| RStudio | Posit PBC, Boston, MA, USA | Version 2023.12.1 |
| Analyst | SCIEX | Version 1.6.8 |
| **Other** |  |  |
| *Ablation laser (NIRL)* | Opotek | *Opolette SE 2731* |
| *Telesto II (OCT)* | Thorlabs | *TEL221PSC2-SP1* |
| *QTRAP 5500* | SCIEX |  |
| *ThermoMixer* | Eppendorf |  |
| *UniVapo* | Eppendorf |  |

### Methods and Protocols

#### Samples

A total of 28 tissue samples originating from the palatine tonsil of 11 patients were included in the study. The samples were collected either during panendoscopy procedures or cancer resections under general anesthesia. After rinsing the samples with 0.9% sodium chloride solution (NaCL), the samples were stored in centrifugation tubes and frozen in liquid nitrogen at −80 °C. Tissue sampling took place in accordance with the World Medical Association Declaration of Helsinki and the guidelines for experimentation with humans by the Chambers of Physicians of the State of Hamburg. All patients have given written informed consent that the removed tissue will be used for research purposes. In accordance with local laws (§12 HmbKHG) and the local ethics committee (Ethics commission Hamburg WF-049/09), The Hamburg Commissioner for Data Protection and Freedom of Information (HmbBfDI) was notified of the collection of head and neck tumor tissue in the context of a biobank. The tissue samples were H&E-stained by the Institute of Pathology for histopathological confirmation, following the standard operating procedures. The diagnosis was independently and blindly confirmed by an expert pathologist.

### Conventional Chamber Ablation Setup

All 28 samples were ablated in an already established ablation setup, that has been described in detail in our previous publication (Stadlhofer *et al*, 2023). The divergent beam emitted by the pulsed nanosecond infrared laser system (Opolette SE 2731, Opotek, Carlsbad, CA, USA) passes through a telescope with two plano-convex lenses (ISP-PX-25-150 and ISP-PX-25-100, ISP Optics Latvia, Riga, Latvia) that collimates the beam. After collimation, the beam is focused by a 150 mm focusing lens (ISP-PX-25-150, ISP Optics Latvia, Riga, Latvia) with a spot diameter of about 150 µm, to achieve a long focal spot with equal energy output. Transverse scanning was performed by a dual-axis scanning mirror (OIM202, Optics in Motion, Long Beach, CA, USA) that was controlled by a data acquisition input/output applicator (USB-6343, National Instruments, Austin, TX, USA). To achieve the maximum repetition rate of 20 Hz, the laser triggering was synchronized to the scanning mirror and timed. A camera path was integrated to enable precise targeting of the sample. Ablation was performed inside a closed ablation chamber with a glass window on the top, to avoid contamination. During ablation the sample was placed on a cooling stage. A membrane pump (Mz 2c Vario, Vacuubrand, Wertheim, Germany) established an airstream from the inlet, where an air filter was placed, to the outlet, where the emerging aerosol was caught on a 10 mm-diameter glass fiber filter (GF50 grade, glass fiber filter without binders, Hahnemühle FineArt, Dassel, Germany). After each ablation, the filter was transferred to a tube for subsequent lipid and protein extraction.

### Handheld Ablation Setup

In total six samples, that were already ablated with the chamber setup, were ablated a second time with the new designed handheld applicator. The divergent beam was emitted by the same pulsed nanosecond infrared laser system (Opolette SE 2731, Opotek, Carlsbad, CA, USA). It passes through a Galilei telescope with one plano-concave lens and one plano-convex lens (ISP-PC-25-75 and ISP-PX-25-150, ISP Optics Latvia, Riga, Latvia) to double the width of the beam and to collimate it. After that, the beam is focused by a 100 mm focusing lens (ISP-PX-25-100) to hit the fiber port optimally. The beam is then transmitted by a 2 m hollow silica waveguide (optimized for 2.94 µm; I.D. bore: 500 µm, Laser Components, Olching, Germany). The end of the laser fiber is mounted on a fiber port on a round handpiece with a diameter of 2 inch that contains a 100 mm plano-convex lens for collimating the beam (ISP-PX-25-100) and a 75 mm plano-convex lens (ISP-PX-25-75) for focusing the beam with a long working distance for ablation. The handpiece was mounted on a stand to ensure an optimal and stable ablation process. The sample was placed beneath the handheld device on a custom-built cooling plate (−12°C), which is mounted on a motorized 2-axis stage. The two stages (MLT25, Newport, Irvine, CA, USA) are synchronized with the laser utilizing the motor controller (XPS-RLD4, Newport, Irvine, CA, USA) to imitate a future scanning in the handheld applicator. A simple webcam was directed at the sample to monitor the ablation process. Pumping and aerosol collection using glass fiber filters were performed identically to the chamber setup.

### Ablation Parameters and Tissue Sampling

In both ablation setups, the NIRL was emitting with a pulse width of 7 ns at 2940 nm to match the O-H vibrational stretching band of water.

The chamber setup was equipped with a scanning mirror. We utilized this advantage for gathering a representative overview of the sample’s molecular composition. Each of the three technical replicates was taken by sampling from four randomly chosen locations (P1-P4), which were pooled on the glass fiber filter and analyzed together. The ablation pattern for each location consisted of 5×5 laser shots with 100 µm spacing, which was applied with 13 repetitions. As reported before, each laser shot measured approximately 100 µm and removed about 25 µm of tissue with a pulse energy of 1.45 mJ at the sample position. This pattern resulted in conical ablations, the volume of which we measured for reference using optical coherence tomography (OCT) imaging.

The handheld setup does not provide scanning; therefore, we used a motorized 2-axis stage, which was synchronized with the laser to apply only one pattern of 10×10 laser shots with a 100 µm spacing and 19 repetitions for each of the two technical replicates. Here, we increased the applied laser shots to compensate for the reduced pulse energy of only 0.7 mJ at sample position, due to losses introduced by the optical elements and the hollow waveguide.

During ablation in both setups, the temperature of the frozen tissue sample was maintained at −10 °C to ensure optimal tissue ablation. The emerging aerosol was guided onto the glass fiber filter and afterwards transferred into a tube and stored until further lipid and protein extraction. Every sample was ablated three times according to the above-described scheme (technical replicates). When the tissue sample was replaced, the filter mount was cleaned with isopropanol as well as with an ultrasonic cleaner (USC100TH, VWR, Darmstadt, Germany) for 5 min. Additionally, during the chamber setup, the inside of the chamber was cleaned with isopropanol.

Histological analysis of the ablated tissue samples by an expert pathologist confirmed their classification as either healthy or OPSCC. The histological images demonstrate the precise cutting capability of the NIRL both within the chamber setup (Figure 11 C) and with the designed handheld applicator (Figure 11 E). The tissue was removed with minimal damage to the adjacent tissue by both approaches. (Figure 11 C, D, E, and F).

**Figure 10.**
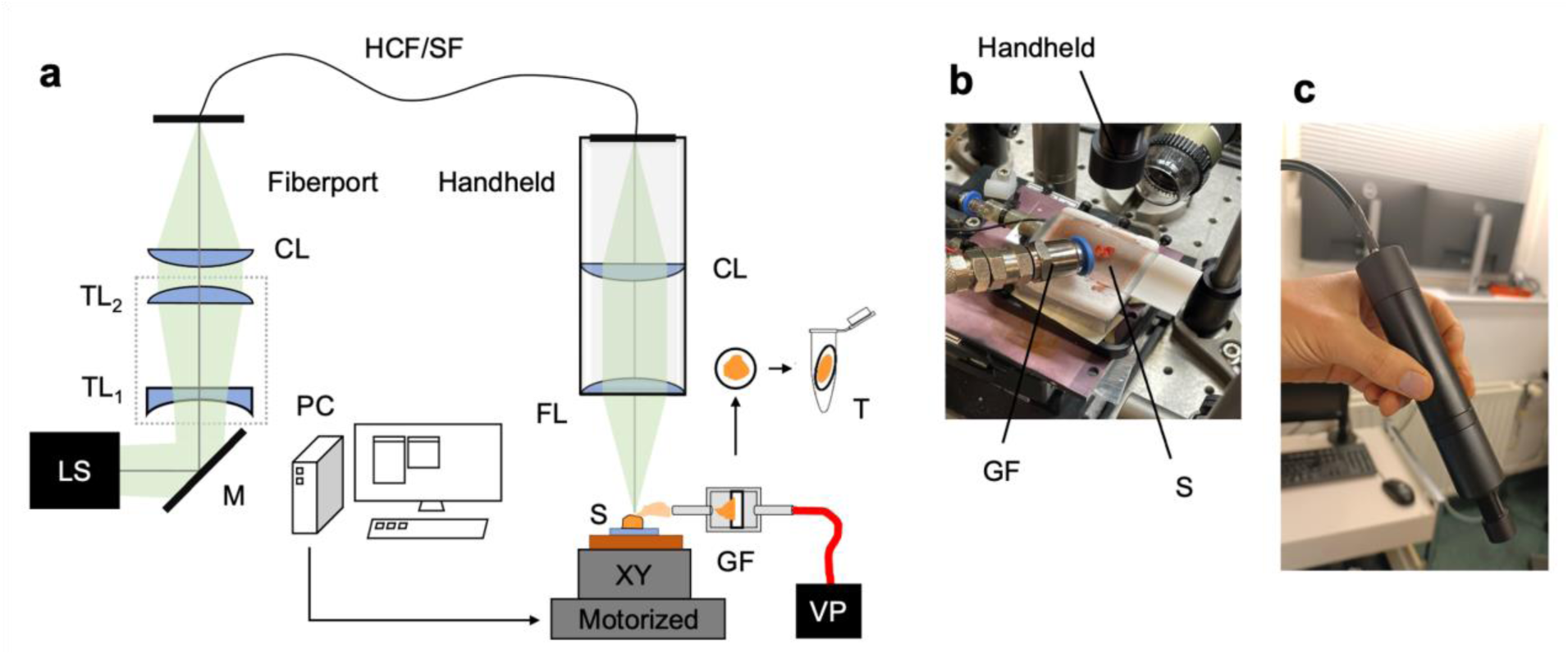
**A** Scheme of the ablation setup. **B** Picture of sample placed below the handpiece. **C** Glass fiber filter after ablation to be placed in the tube. LS: nanosecond infrared laser system; M: mirror; TL1/2: telescope lens 1/2; GT: Galilei telescope; CL: collecting/collimating lense LF: laser fiber; PC: computer; XY: motorized 2-axis stage; S: frozen sample; GF: glass fiber filter; VP: vacuum pump; T: reaction tube

**Figure 11.**
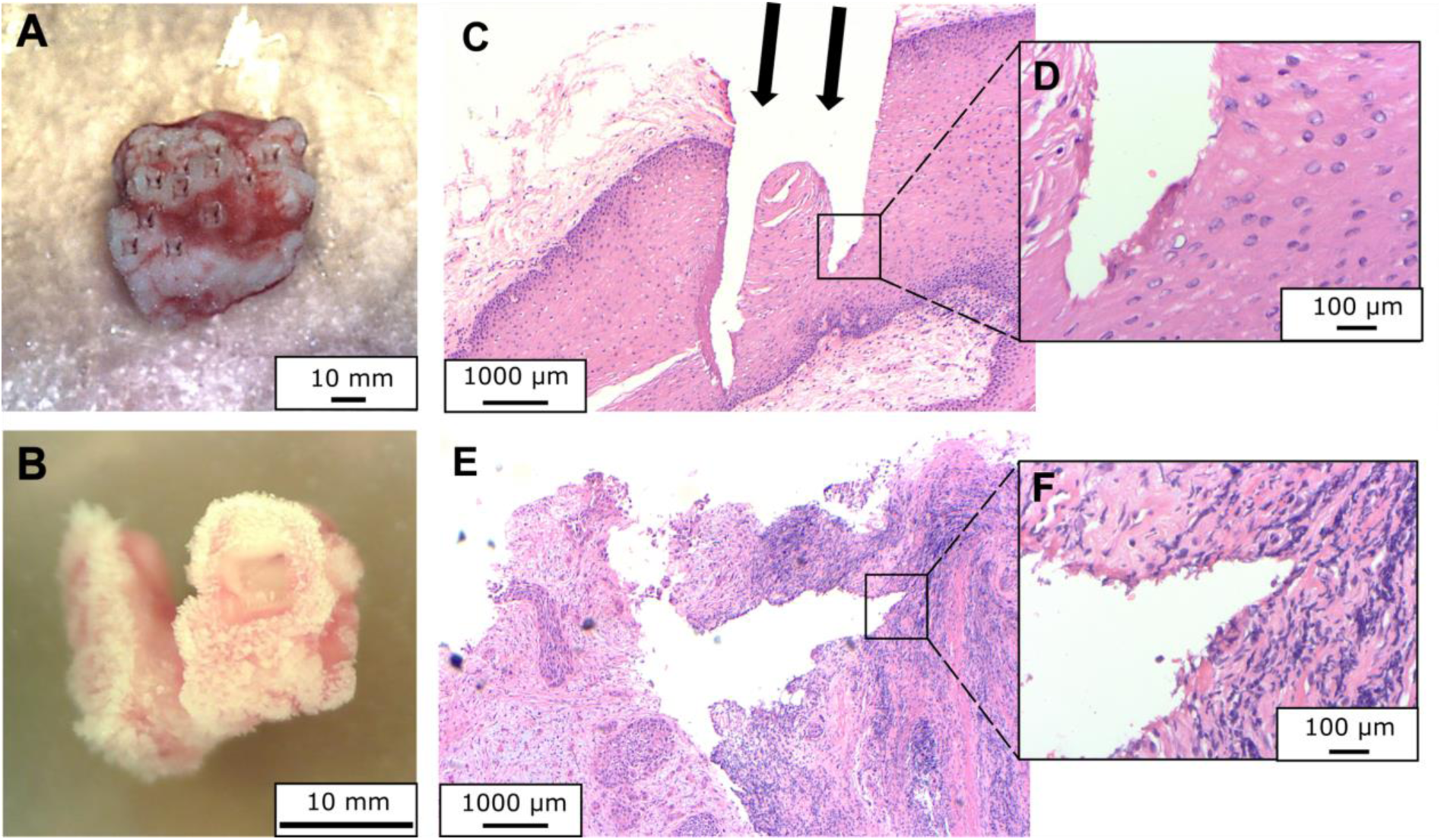
**A** Sample in the chamber after ablation **B** Sample after ablation with handheld device **C** Histological H&E stained picture of a healthy sample after ablation with chamber setup **D** Low damage applied to the surrounding tissue using NIRL **E** Histological H&E stained picture of a healthy sample after ablation with handheld applicator **F** Low damage applied to the surrounding tissue using NIRL with handheld applicator.

### Determination of Ablation Volume with Optical Coherence Tomography

The sample surface with the ablations in the chamber setup was measured using an optical coherence tomography imaging system with a center wavelength of 1300 nm (TEL221PSC2-SP1, Thorlabs, Lübeck, Germany). Based on the 3D OCT image data (Figure 12 A-C), which were processed in the opensource software Fiji (Schindelin *et al*, 2012), manual segmentation was performed using the opensource software ITK-Snap (Yushkevich *et al*, 2006) to determine the ablation volume of the locations (Figure 12 D, top). The segmented label was interpolated for volumetric quantification, resulting in an ablation volume 45 nL per location (Figure 12D, bottom) and a total of 180 nL per replicate.

**Figure 12.**
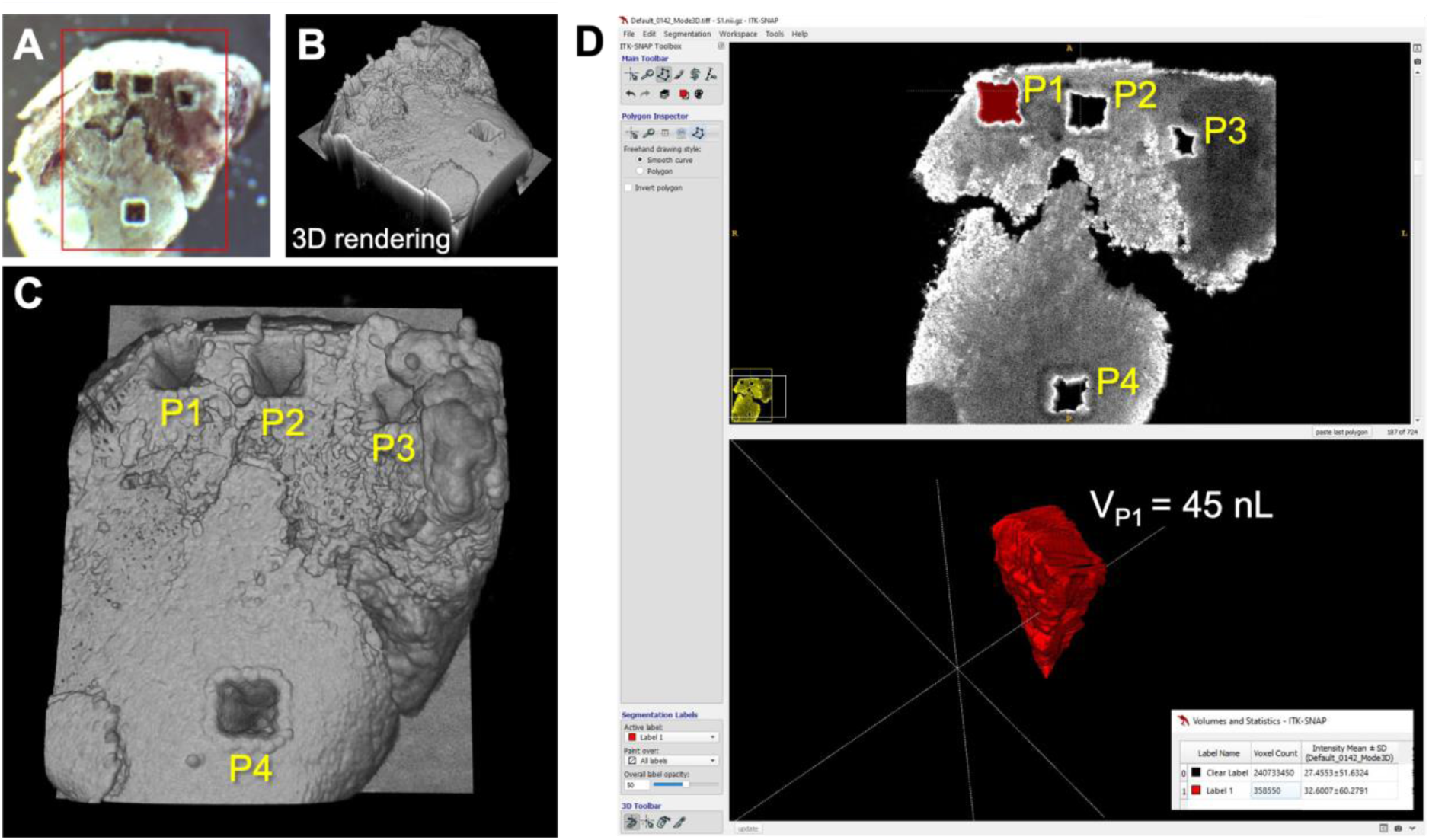
**A** Camera view of the OCT system of an ablated sample from the chamber setup. **B** 3D rendering of the acquired OCT image data. **C** View of the four ablation locations (P1-P4) forming one of the three replicates for each ablated sample. **D** Graphical user interface of the opensource software ITK-Snap for manual segmentation of the location P1 (top) with volumetric quantification (bottom).

### Lipid Extraction

Before lipid extraction, a mixture of deuterated internal standards was added to the samples according to Table 4.

**Table 4.** Components of internal Standard mix.

|  | Avanti # | Final amount /sample (µg) |
| --- | --- | --- |
| Ultimate SplashOne | 330820 | Various |
| dFA 18:1 | 861809 | 0.1 |
| dCer d18:0/13:0 | 330726 | 0.01 |
| Glu Cer(d18:1-d7/15:0) | 330729 | 0.2 |
| dLacCER d18:1/15:0 | 330727 | 0.1 |
| 15:0-18:1-d7-PA | 791642 | 0.1 |

Following the addition of 25 µL of the internal standard solution, 500 µL of MTBE/MeOH (3:1, v/v) were added to the glass fiber filter for the lipid extraction. The samples were vortexed for 30 seconds and then incubated for a further 15 minutes at 4 °C. Subsequently, 325 µL of MeOH/H2O (3:1, v/v) were added to the samples, which were then vortexed for 30 seconds. The samples were subjected to centrifugation for 5 minutes at 4 °C with an acceleration of 20,000 g. From each sample, 300 µL of the upper lipid-containing phase were transferred into new reaction tubes. The lipid phase samples were then subjected to vacuum evaporation.

### Lipidomic Analysis

Tissue samples were analyzed using a shotgun lipidomics approach as described by Su et al. (Su *et al*, 2021). Prior to mass spectrometric measurements, the samples were resuspended in 275 µL of the sample running solution, which consisted of 10 mM ammonium acetate in DCM/MeOH (1:1, v/v).

The reconstituted lipid extracts were directly infused into the mass spectrometer using an ultrahigh-pressure liquid chromatography system (Nexera X2, Shimadzu, Kyoto, Japan) with dichloromethane (50): methanol (50) containing 10mM ammonium acetate as eluent at a flowrate of 0.008 mL/min. Ionization was achieved using a Turbo V source with a 65 μm ESI Electrode. Targeted lipid analysis was conducted by a QTRAP® system (QTRAP® 5500; SCIEX) run in multiple reaction monitoring mode with polarity switching operated via Analyst (version 1.6.8, SCIEX). In particular, samples were measured twice, using two different MRM methods. Method 1 contained a lipid separation step via differential mobility spectrometry employing the SelexION technology after ionization to quantify PC, PE, PG, PI, PS, and SM and method 2 was employed to quantify CE, Cer d18:0, Cer d18:1, DG, FFA, HexCER, LPC, LPE, LacCER, and PA. Method details and MRM transitions can be downloaded from github (https://github.com/syjgino/SLA). SLAv1.3 and in particular keyV4 were used. After acquisition, MSconvertGUI (Version 3.0.21245-5724be1) was employed to convert the raw data obtained from the mass spectrometer into mzML format. Further data processing and lipid quantification was enabled by using the Shotgun Lipidomic Assistant (SLA) software, a python-based application according to Su et al. (Su *et al*, 2021).

### Data Analysis and Visualization

Quantified lipid species concentrations of the chamber setup were uploaded to RStudio (version 2023.12.1+402, Posit PBC, Boston, MA, USA). The data were log_2_ transformed. Lipid species were filtered to retain only those with at least 70% valid values per species. Afterwards, missing values were replaced using kNN imputation (k=5). Welch’s test (p-value ≤ 0.05, two-fold change) was performed on each patient’s corresponding healthy and tumorous tissue. oPLS-DA was conducted using MetaboAnalyst.ca. Quantified lipid species concentrations of the handheld setup were analyzed separately using the same workflow.

## Acknowledgements

This work was supported by a DFG grant (SCHL 406/21-1) in the funding program: “New instruments for research” and by the Mildred Scheel Cancer Career Center Hamburg (HaTriCS4).

## Author Contribution

JH, DE, MM and LK conceived the study and designed all experiments.

Ablation setup and handheld design and OCT measurements by JH.

LK carried out the ablation experiments with supervision of JH.

LK, MM, JH, AW analyzed the lipid data and JH the imaging data.

LK and MM performed the statistical analysis.

HZ collected and categorized the samples.

TSC and WW performed the histological analysis and the provided images.

Supervision and administration were done by JH and AB.

AW, MM and JHe established the MS methods and performed the shotgun lipidomics measurements.

AB, DE, HS assisted in the design, analysis, and interpretation of experiments.

Resources were provided by JHe, AB, DE, CSB, JH and HS.

All authors discussed the results and LK, MM, and JH wrote the original draft of the manuscript. Review and editing were done by JH, MM, HS, CSB, TSC, AW, JHe, DE and AB.

## The Paper Explained

### Problem

Oropharyngeal Squamous Cell Carcinoma (OPSCC) is a life-threatening cancer with rising incidence over the past decade. As Surgery is one mainstay of therapy, the intraoperative assessment of tumor margins needs to be improved. Nanosecond infrared lasers (NIRL) enable the fast extraction of intact biomolecules, such as lipids, from tissue. Subsequent mass spectrometric (MS) analysis may allow for the rapid differentiation between tumorous and healthy tissue. However, there is currently a lack of knowledge regarding marker lipids for OPSCC.

### Results

Our study reveals lipidome differences in the OPSCC tissue, despite biological heterogeneity of the samples. We identified potential biomarkers suitable for future targeted lipidomic analysis; however, these findings require further validation in larger patient cohorts. Enhancing the clinical applicability, we developed a custom handheld applicator capable of performing tissue ablation without thermal damage. The lipidome differences observed were detected using the handheld applicator as well.

### Impact

The study creates a base for further studies, as not the whole lipidome needs to be analyzed but only a small number of significantly altered lipid species. It proves that a handheld NIRL coupled to MS is a realistic and applicable option for future applications on patients.

